# Dissection of neuropsychiatric-relevant behavioral and structural phenotypes utilizing neuron type-specific ablation of the *Bassoon* gene in mice

**DOI:** 10.64898/2026.10.09.757973

**Authors:** Judith C. Kreutzmann, Frank Angenstein, Anna Fejtova, Oliver Stork, Eckart D. Gundelfinger, Markus Fendt, Anil Annamneedi

## Abstract

Rare pathogenic mutations in the human *BSN* gene, which encodes the presynaptic cytomatrix protein Bassoon – a key organizer of the neurotransmitter release apparatus – have been linked to neuropsychiatric conditions including schizophrenia. Here we began to dissect the potential contribution of *Bsn* deficiency in distinct populations of mouse (fore)brain neurons on behavioral and anatomical alterations resembling schizophrenia-like phenotypes.

To this end, conditional knockout mice lacking Bassoon in subsets of inhibitory synapses (*Bsn^Dlx5/6^*cKO), in forebrain excitatory synapses (*Bsn^Emx1^*cKO), or at dopaminergic release sites (*Bsn^DAT^*cKO) were generated. The mice were assessed using prepulse inhibition, a paradigm that models sensorimotor gating, which is often impaired in schizophrenia, and a fear/safety learning paradigm. Further, anxiety-related behaviors were evaluated and magnetic resonance imaging (MRI) was performed to assess anatomical brain alterations.

Our data reveal synapse-specific effects of the Bassoon deficiency. Sensory gating deficits were observed in both *Bsn^Dlx5/6^*cKO and *Bsn^Emx1^*cKO mice. *Bsn^Dlx5/6^*cKO mice further demonstrated impaired discrimination between conditioned fear and safety – a condition frequently observed in individuals with schizophrenia. By contrast, *Bsn^DAT^*cKO mice displayed enhanced discrimination between conditioned fear and safety. All mouse strains showed elevated anxiety-like behavior in the light/dark box. Structural MRI revealed increased brain volumes in *Bsn^Dlx5/6^*cKO and *Bsn^Emx1^*cKO mice, with the most pronounced changes in the hippocampus and cortex.

Previous studies have shown that *Bsn^Dlx5/6^*cKO mice display autistic-like behaviors, altered hippocampal network activity and epilepsy. Our findings support the notion that deficits in *Bsn*-mediated functions, particularly in interneurons, may contribute to the etiology of neurodevelopmental and neuropsychiatric conditions.

**Highlights:**

- Loss of Bassoon in inhibitory or excitatory synapses induces macrocephaly and prepulse inhibition deficits.
- Selective ablation of *Bsn* in inhibitory or dopaminergic neurons exerts opposite effects on fear/safety learning.
- All three conditional *Bsn* mutants – lacking Bassoon in inhibitory, excitatory or dopaminergic neurons – exhibit an anxiety-like phenotype.

**Graphical abstract:** 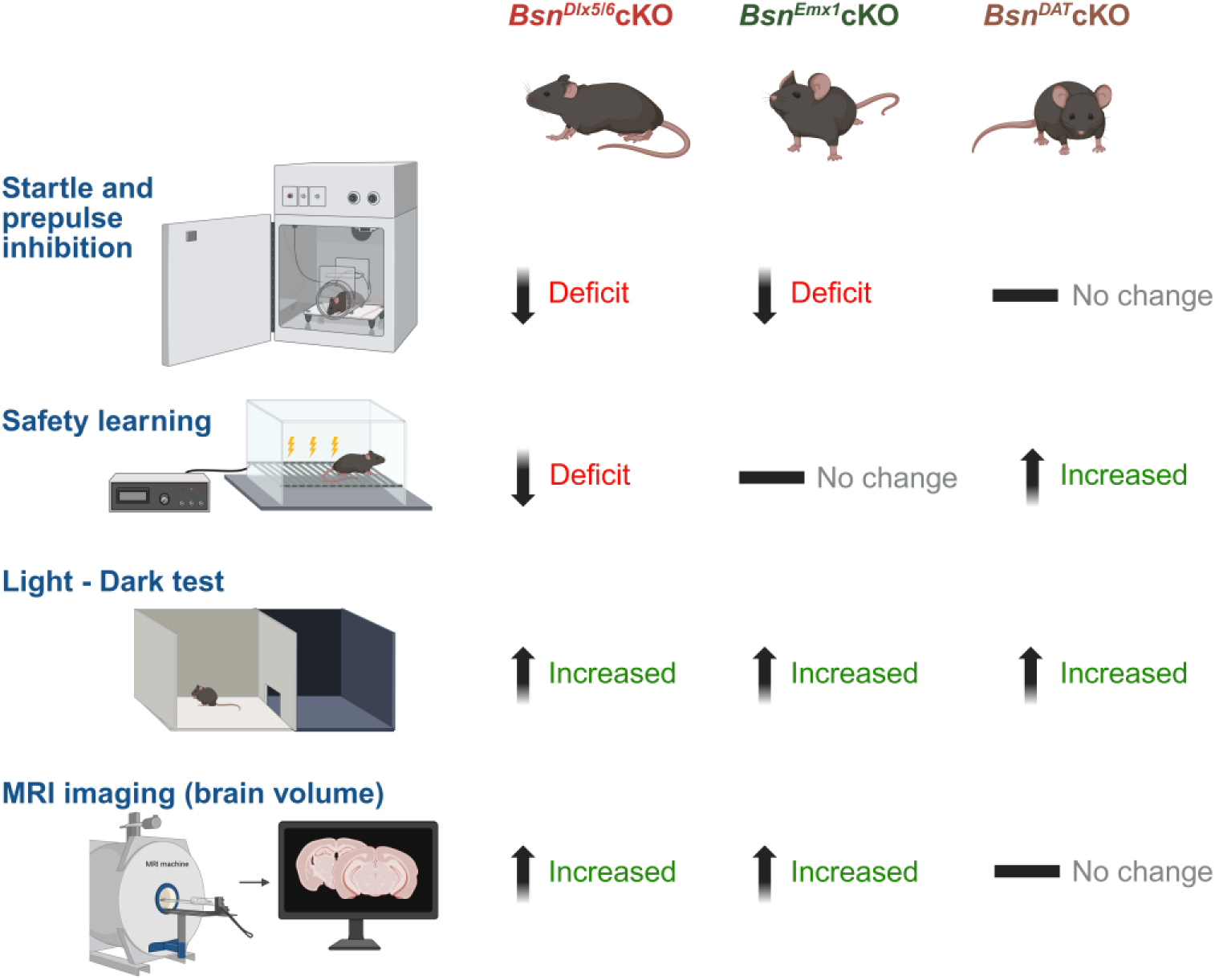

## INTRODUCTION

Memory formation depends on the coordinated activity of distinct brain regions, where learned behaviors are encoded through changes in signal processing within brain neuronal networks. Changes in the synaptic transmission between neurons linked to activity-dependent changes in synaptic protein composition importantly contribute to this process. One of the key synaptic proteins is Bassoon, a large (420 kDa), multidomain scaffolding protein encoded by the *BSN* gene in humans (*Bsn* in mice). Located in the presynaptic cytomatrix at the active zone (CAZ), Bassoon plays a critical role in the assembly and plasticity of the presynaptic neurotransmitter release apparatus. Bassoon along with its interaction partners contributes to the organization of the synaptic vesicle cycle at the active zone, phosphorylation at presynapse and also contributes to the activity-dependent regulation of neuroplasticity genes via controlling presynapse-to-nucleus signaling (Ackermann et al., 2015; Gundelfinger et al., 2016; Gundelfinger and Fejtova, 2012; Ivanova et al., 2015; Montenegro-Venegas et al., 2022; Südhof, 2012; Ziv and Garner, 2004).

To date, few studies have investigated the systemic role of Bassoon. *Bsn* mutant mice lacking the central part of the protein (*Bsn*^ΔEx4/5^; (Altrock et al., 2003) have been reported to exhibit a range of behavioral phenotypes, including epileptic seizures, impaired performance in socially transmitted food preference task, and enhanced performance in a two-way active avoidance paradigm (Blondiaux et al., 2023; Ghiglieri et al., 2010; Sgobio et al., 2010). Mice lacking *Bsn* specifically in a subset of GABAergic interneurons developmentally controlled via the *distal-less Dlx5/6* regulatory element (Wang et al., 2010; Zerucha et al., 2000) (*Bsn^Dlx5/6^*cKO) develop autism-like behavioral and physiological phenotypes, associated with disturbed brain metabolism (Annamneedi et al., 2026) and epilepsy (Blondiaux et al., 2023). Furthermore, we previously demonstrated that mice with a conditional knockout of *Bsn* in excitatory neurons of the forebrain (*Bsn^Emx1^*cKO) show enhanced dentate gyrus (DG)-dependent contextual fear memory and improved spatial pattern separation learning, which appear to result from disturbed maturation of the DG (Annamneedi et al., 2018). Interestingly, as opposed to *Bsn^Dlx5/6^*cKO mice, *Bsn^Emx1^*cKO mice display enhanced performance in an auditory cortex-dependent frequency-modulated tone discrimination (FMTD) task (Oelschlegel et al., 2025). Moreover, mice lacking *Bsn* in neuron of telencephalic excitatory circuits (*Bsn^NEX^*cKO) have been reported to display enhanced memory retention, however, at the expense of cognitive flexibility (Zoicas et al., 2026).

Symptoms of schizophrenia include deficits in sensorimotor gating, enhanced anxiety, impaired discrimination learning, and disturbances in the assignment of stimulus relevance (Braff and Geyer, 1990; Hofer et al., 2001; Holt et al., 2012, 2009; Lubow and Weiner, 2010; Lubow, 2005; Temmingh and Stein, 2015). Among these features, impairments in safety learning -the ability to correctly identify stimuli that predict the absence of threat and to discriminate them from stimuli predicting threat-are consistently observed (Lubow and Weiner, 2010). Such safety learning deficits are often assessed using fear/safety discrimination paradigms (Qiao et al., 2025).

Early postmortem studies revealed elevated Bassoon expression in the prefrontal cortex of individuals with schizophrenia, a brain region critically involved in the disorder’s cognitive and emotional deficits (Martins-de-Souza et al., 2009). Other studies have demonstrated an interaction between Bassoon and the schizophrenia susceptibility gene D-amino acid oxidase (DAO), suggesting a role in modulating glutamatergic signaling (Popiolek et al., 2011). More recently, a rare pathogenic mutation in Bassoon has been associated with neurodevelopmental and neuropsychiatric conditions (Chen et al., 2021; Guzman et al., 2025), suggesting that altered Bassoon function may contribute to synaptic dysfunction observed in these disorders. Together, these findings indicate that Bassoon-dependent presynaptic mechanisms may be relevant for neural circuits involved in stimulus relevance processing and safety learning, processes that are disrupted in schizophrenia (Lisman et al., 2008; Uhlhaas and Singer, 2010). Importantly, safety learning and sensorimotor gating depend on the interaction of excitatory, inhibitory, and dopaminergic neurotransmission (Gray, 1998; Lewis et al., 2005; Lodge and Grace, 2011). Thus, neuron type-specific manipulation of presynaptic scaffolding proteins such as Bassoon provides a strategy to dissect circuit mechanisms underlying schizophrenia-relevant learning alterations with high cellular specificity.

The aim of the present study was to investigate whether the loss of the pre-synaptic scaffolding Bassoon in specific neuronal populations leads to behavioral alterations relevant to neuropsychiatric disorders, with particular focus on schizophrenia-related phenotypes. To address this, we established and used conditional knockout mouse lines lacking Bassoon specifically in a subset of inhibitory synapses (*Bsn^Dlx5/6^*cKO) (Annamneedi et al., 2026), in forebrain excitatory synapses (*Bsn^Emx1^*cKO) (Annamneedi et al., 2018) and established conditional knockout mice lacking Bassoon in dopaminergic synapses (*Bsn^DAT^*cKO). The mice were subjected to a comprehensive behavioral assessment, including paradigms linked to core features of schizophrenia, such as prepulse inhibition (PPI) of the acoustic startle response (ASR), which evaluates sensorimotor gating. To directly assess safety learning, mice were tested using a fear/safety learning paradigm conceptually related to latent inhibition, modelling impaired stimulus relevance assignment commonly reported in schizophrenia (Holt et al., 2012; Lubow, 2005). Additional tests, such as the light/dark box for anxiety-like behavior and brain volumetric analysis via MRI, were employed to further characterize relevant consequences of neuron type-specific loss of Bassoon expression.

## MATERIAL AND METHODS

### Animals and Housing Conditions

Experimental subjects were adult 8-14 weeks-old male and naturally cycling female conditional knock-out mice and their wild-type littermates. All mice were bred in the animal facility of the Leibniz Institute for Neurobiology (LIN) in Magdeburg. Before and after behavioral testing, mice were housed and cared for at the Institute for Pharmacology and Toxicology (IPT) and the Institute of Biology (IBIO), Otto-von-Guericke University (OvGU) Magdeburg. At all sites, mice were group-housed by sex (4-6 littermates/cage) in transparent Makrolon Type III cages (840 cm^2^) or GM-500 seal safe plus cages (Techniplast) with wood chip bedding, nesting material and cage enrichment. The animals had free access to standard chow (Ssniff^®^ R/M-H, V1534-0, Germany) and tap water, with a fixed 12:12 hour light/dark photoperiod (lights on at 06:00 h) in a temperature-(22 ± 2 °C) and humidity-controlled (55 ± 10 %) room.

All experimental procedures were approved by the local authorities (Landesverwaltungsamt Sachsen-Anhalt: Az. 42502-2-988 LIN, 42502-2-1303 LIN and 42502-2-1309 Uni MD) and conducted in agreement with international guidelines and regulations for animal experiments (2010/63/EU).

### Cell type-specific *Bassoon* conditional Knock-Out (cKO) Mice

Three different conditional *Bsn* knockout mice were used in this study including *Bsn^Dlx5/6^*cKO – lacking *Bsn* in nearly 40% of inhibitory interneurons (Annamneedi et al., 2026; Blondiaux et al., 2023), *Bsn^Emx1^*cKO – lacking *Bsn* in forebrain excitatory neurons (Annamneedi et al., 2018) and *Bsn^DAT^*cKO – lacking *Bsn* in dopaminergic neurons. Generation of *Bsn^DAT^*cKO mice was performed applying a strategy analogous to the previously reported generation of *Bsn^Emx1^*cKO (Annamneedi et al., 2018). Briefly, *Bsn^lxlx^* (B6.Cg*-Bsn^tm1.1Arte^*; (Annamneedi et al., 2018) mice were bred with mice expressing Cre recombinase under DAT (Dopamine transporter) promoter (*DAT^IRES-^* ^Cre^ mice: B6.SJL-*Slc6a3^tm1.1(cre)Bkmn^*^/J^, The Jackson Laboratory; (Bäckman et al., 2006). Cre-mediated recombination resulted in deletion of the N-terminal region of the Bassoon protein, leading to a frameshift and the introduction of a premature stop codon in the target cell populations. All mouse lines including *Bsn^lxlx^* and *DAT^IRES-Cre^* were maintained on a C57BL/6NCrl background. *Bsn^lx/lx^* mice lacking Cre expression served as controls for the *Bsn^Dlx5/6^* and *Bsn^Emx1^* lines, whereas *Bsn*^+/+^ X *DAT^IRES-^*^Cre^ mice were used as controls for the *Bsn^DAT^* line.

Genotyping of *Bsn^lxlx^*, *Bsn^Emx1^*cKO and *Bsn^Dlx5/6^*cKO mice was performed as described previously (Annamneedi et al., 2026, 2018). For *DAT^IRES-^*^Cre^ mice, genotyping was performed with slight modifications to the protocol provided by The Jackson Laboratory. PCR amplification was carried out using Qiagen Taq polymerase (Qiagen, catalog#201207). The thermal cycling conditions were as follows: initial denaturation at 94 °C for 3 minutes; 35 cycles of 94 °C for 30 seconds, 62 °C for 30 seconds, and 72 °C for 30 seconds; followed by a final extension at 72 °C for 5 minutes.

### Immunohistochemical Analyses

Immunohistochemical procedures were performed as described previously (Annamneedi et al., 2018). 30 µm striatal coronal sections were used to analyze the expression of Bassoon in dopaminergic terminals. Primary antibodies included mouse anti-Bassoon; (Enzo Life Sciences Inc. New York, USA; RRID: 1:1000) and rat anti-DAT (Abcam, Cambridge, UK; RRID: 1:500). Secondary antibodies used were goat anti-mouse Cy3 (Jackson Immuno Research Labs, Pennsylvania, USA; RRID: 1:500) and goat anti-rat Alexa 488 (Invitrogen, California, USA; RRID: 1:200). Confocal images were acquired as Z-stacks with a 0.5 μm Z-step size (∼ 5 μm Z-stack volume) using a Leica SP5 confocal microscope equipped with a 63× objective and LCS software (Leica Microsystems, Wetzlar, Germany).

### Behavioral Testing

Behavioral characterization was performed across different laboratories depending on the paradigm employed. Testing of the ASR, PPI and conditioned safety learning was performed during the light phase at the IPT, OvGU, Magdeburg. For these experiments, male and female cKO mice, as well as their wildtype littermates were used: *Bsn^lxlx^* (n = 10 males and n=10 females) and *Bsn^Dlx5/6^*cKO (n = 12 males and n = 7 females); *Bsn^lxlx^* (n = 14 males and n = 12 females) and *Bsn^Emx1^*cKO (n = 29, 14 males); *DAT^Cre/+^* (n = 12 males and n = 9 females) and *Bsn^DAT^*cKO (n = 9 males and n = 11 females). Although a main effect of sex was observed for the ASR magnitude (males exhibit larger responses), this effect can be attributed to physiological factors, as males generally possess greater body weight/muscle mass and therefore exhibit a more intense startle response. Apart from ASR, no significant sex differences were detected; thus, data from male and female mice were pooled for behavioral analysis. General behavioral characterization of male *Bsn^DAT^*cKO mice (see Supplementary information) was conducted at the IBIO, OvGU, Magdeburg.

Mice were transferred to the respective laboratories at least 7 to 10 days prior to start of the experiments to allow for habituation to the new environment. Mice were maintained under the same housing and light conditions as described above, with the exception of those used for the general characterization of *Bsn^DAT^*cKO mice. These animals were single-housed immediately upon transfer and kept under a reverse light-dark cycle (12 h light/12 h dark cycle, lights on at 7 p.m.), at a constant room temperature of 22 ± 2 °C. Behavioral characterization for *Bsn^DAT^*cKO mice was performed during the dark phase and included a comprehensive test battery, including home cage activity monitoring, open field, sucrose drinking test, inverted grip strength test, rotarod, and fear conditioning (see Supplementary information).

#### Acoustic Startle Response and Prepulse Inhibition

##### Setup

A computerized startle system (SR-LAB, San Diego Instruments, USA) with eight chambers (35 cm × 35 cm × 38 cm) was used. Each chamber was equipped with a platform with a transparent horizontal cylinder-shaped enclosure for the animal (10 cm x 4 cm) and a high-frequency loudspeaker for delivering the acoustic stimuli. A piezoelectric motion sensor was mounted below the platform to measure the animals’ movement. During each test session, a background noise with an intensity of 50 dB SPL was presented to mask environmental noises. The delivery of all stimuli was controlled by the SR-LAB software.

##### Acoustic Startle Response

To measure the startle magnitude to acoustic stimuli with different intensities, mice were placed into the enclosures and after a 5-minute acclimation period, 24 startle stimuli of different intensities (78, 84, 90, 96, 102, 108, 114, 120 dB SPL) were presented pseudo-randomly with an inter-trial interval (ITI) of 20 s.

##### Prepulse Inhibition

Following 5 minutes of acclimatization (background noise only), a baseline measurement of 12 startle stimuli (white noise, 40 ms, 108 dB SPL) with an ITI of 20 s was obtained. Following the baseline, 6 blocks each including a startle stimulus alone or startle stimuli with prepulses of 2, 4, 8, 12, or 16 dB SPL above background noise (60 dB SPL) were presented in a pseudo-randomized order. All prepulses had a duration of 20 ms and preceded the startling stimulus by 100 ms (onset to onset).

#### Light/Dark Box Test

To investigate anxiety, we used a computerized light/dark box system (TSE Systems, Bad Homburg, Germany) described in detail elsewhere (Kreutzmann et al., 2024). On the test day, the mice were placed into the middle of the dark compartment and allowed to explore both compartments for five minutes. Movements of the animals were detected by infrared sensors (distance: 14 mm) and analyzed by TSE Phenomaster software (Version 4.9.4). As a behavioral readout for trait anxiety, we primarily used the percent time and distance spent in the light compartment.

#### Conditioned Fear/Safety Learning

##### Setup

For the conditioned fear/safety learning procedure, we used a computerized fear-conditioning system (TSE Systems, Bad Homburg, Germany), consisting of four transparent Perspex box (46 cm × 46 cm × 32 cm) that were surrounded by infrared animal detection sensor frames. The boxes were in sound-attenuating chambers equipped with loud-speakers for acoustic stimuli, a light source (continuous background illumination of ∼20 lux) and ventilation fans, producing a background noise of approximately 55 dB SPL. The floors of the boxes consisted of removable stainless-steel grids (bars: 4 mm in diameter with 9 mm between the bars) that were connected to a shock unit and able to deliver electric stimuli. Delivery of all stimuli was controlled by the TSE FC Software (V9.10). To ensure that mice received the electric stimuli, animals were monitored via video cameras and the locomotive response to these stimuli was measured for each animal. Movements of the animals were detected by infrared sensors (distance: 14 mm). Freezing behavior was defined as no infrared beam crosses for more than 1 s.

##### Conditioned Fear/Safety Learning

Behavioral experiments were performed using a fear/safety conditioning protocol described previously (Kreutzmann et al., 2020b). The protocol was designed to assess the ability of mice to discriminate between a tone stimulus predicting safety and a context associated with threat.

On day one, mice were habituated to the conditioning boxes and received a single presentation of the tone (10 kHz, 85 dB, 30 s). On days 2 and 3, mice underwent safety conditioning. Each session consisted of five presentations of the tone (conditioned stimulus: CS: 30 s), explicitly unpaired with a scrambled electric stimulus (foot shock: 0.4 mA, 2 s; inter-stimulus interval: 120 s). Thus, the tone was repeatedly presented in the absence of the aversive stimulus and acquired safety-related significance, whereas the conditioning context became associated with occurrence of the aversive stimulus and consequently acquired threat-related significance.

On day four, learned safety was assessed in an expression session. Mice were placed in the conditioning box for 7.5 minutes, during which the safety CS was presented 5 times for 30 s (inter-stimulus interval: 60 s). Freezing behavior was quantified during the 30-s period immediately preceding each CS presentation, reflecting contextual fear, and during each CS presentation, reflecting the learned safety response. Safety discrimination was assessed by comparing freezing during the threat-associated context with freezing during the safety CS. Effective safety learning is therefore reflected by reduced freezing during the CS relative to the surrounding context.

### Anatomical MRI

Mice were anesthetized with 2% isoflurane (in 50:50 N2:O2, v:v) and fixed using a head-holder with bite bars to reduce motion artifacts. After 10 min isoflurane concentration was reduced to 1.2%. MRI experiments were performed on a Bruker BioSpec 94/20 USR scanner at 9.4 T (free bore of 20 cm) equipped with a B-GA12S (440 mT/m) gradient system. A 1H MRI CryoProbeTM 2 element array kit for mice (Bruker BioSpin MRI GmbH, Ettlingen, Germany) was used for signal reception. Heating was provided from the ventral side and heart- and breathing rates were monitored during the entire experiment. Anatomical images were obtained using a RARE (rapid acquisition relaxation enhanced) sequence (Hennig et al., 1986) with the following parameters: TR 3200 ms; effective TE 33 ms, slice thickness 0.6 mm, FOV 20 x 20 mm, matrix 256 x 256 (in plane resolution 78 x 78 µm), RARE factor 8, averages 4. The total scanning time was 6 min 49 s.

### Descriptive and Statistical Analysis

For ASR, the mean startle magnitude for each stimulus intensity was calculated for each individual. For PPI, the prepulse inhibition for each prepulse intensity was calculated for each individual according to the following formula: PPI = (mean startle magnitude without prepulse - mean startle magnitude with prepulse) / (mean startle magnitude without prepulse/100). Furthermore, the mean PPI of the individual animals was calculated, i.e. the mean PPI for all prepulse intensities. For the conditioned fear/safety procedure, the mean percent in freezing to the context and the tone stimulus (later safety CS) were calculated for the acclimation and expression sessions. Furthermore, the percent effect of the safety CS related to contextual freezing for each individual was calculated.

Statistical analyses were performed using Prism 10.0 (GraphPad Software Inc., La Jolla, CA, USA). Statistical significance for the mean PPI magnitude, light/dark box read-outs, as well as the individual percent effect of the safety CS related to contextual freezing and the shock-induced activity was analyzed with Student’s two-tailed t-test. ASR, PPI and percent context freezing and safety CS freezing (fear inhibition) during the acclimation and expression session was analyzed by multi-factorial analyses of variance (ANOVA) with genotype as between-subject factors, and trial type (context, safety CS) or stimulus intensity as within-subjects factor. Between-subjects and within-subject post-hoc comparisons were made using Sidak’s multiple comparisons test. Main effects and interactions were deemed significant with p ≤ 0.05 for all statistical tests. Results are represented as mean ± SEM.

## RESULTS

### Characterization of *Bsn* cKO Mice

Bassoon expression patterns in the mutant mouse strains *Bsn^Emx1^* and *Bsn^Dlx5/6^* have been described previously (Annamneedi et al., 2026, 2018; Blondiaux et al., 2023). While Bassoon cluster localization serves as a key indicator of organization of the dopaminergic synapses during early postnatal development, its specific contribution to dopamine-mediated behavioral functions remains largely unexplored (Cui et al., 2021). We assessed the depletion of Bassoon from dopaminergic release sites in the newly generated *Bsn^DAT^*cKO mice using immunohistochemistry. Sections of the dorsal striatum from control and cKO mice were stained with antibodies against Bassoon and the dopaminergic marker DAT (Figure 1). In control mice, Bassoon and DAT immunoreactivities overlapped, which was absent in *Bsn^DAT^*cKO mice, indicating a loss of Bassoon expression in DAT positive synapses. Similar observations were made for the ventral striatum (Kalinichenko et al., 2026). These results confirm the selective loss of Bassoon from dopaminergic release sites in the *Bsn^DAT^*cKO mouse line.

**Figure 1:**
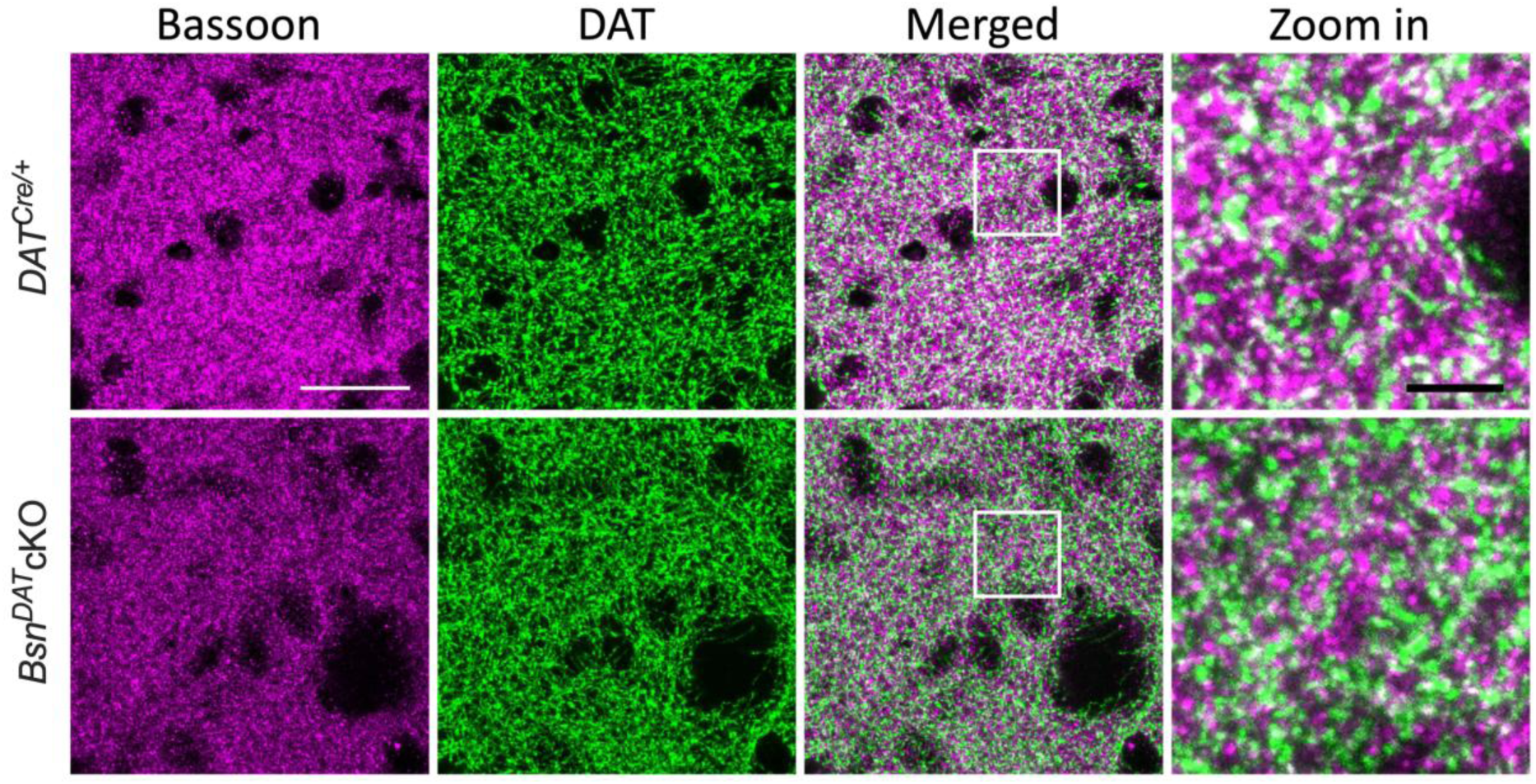
Characterization of *Bsn^DAT^*cKO mice. Immunohistochemical staining of striatal sections from control (*DAT^Cre/+^*) and *Bsn^DAT^*cKO mice using anti-Bassoon and anti-DAT antibodies. Co-localization of Bassoon at dopaminergic synapses is observed in controls as indicated by the merging of magenta and green signals (white colored puncta). By contrast, this overlap was basically absent in *Bsn^DAT^*cKO mice, with the two signals clearly separated, indicating a reduced co-localization in *Bsn^DAT^*cKO mice (merged and zoom in). Scale bar is 25 μm and 5 μm (zoom in).

Total Bassoon levels seemed to be slightly reduced (i.e. by about 10%) in striatal tissue from *Bsn^DAT^*cKO mice, as quantified by immunoblotting of the P2 crude membrane fraction (data not shown). Such a modest reduction is consistent with the proportion of dopaminergic terminals expressing DAT in mouse striatum (Paget-Blanc et al., 2022).

In the general behavioral characterization, *Bsn^DAT^*cKO mice did not display pronounced abnormalities compared to control (*DAT^Cre/+^*) mice (Supplementary Figures 1 and 2). Total activity in home cages was unaltered; although *Bsn^DAT^*cKO mice displayed a modest increase in activity during the dark phase of the circadian cycle (Supplementary Figure 1A-C). In the open-field arena, the novelty-induced activity and anxiety-like behavior were comparable between the groups (Supplementary Figure 1D, E). Similarly, sucrose consumption rate (Supplementary Figure 1F), performance in rotarod test (Supplementary Figure 1G, H) and fear conditioning (Supplementary Figure 1I, J) were unaltered in *Bsn^DAT^*cKO mice compared to controls.

### Bassoon Deficiency Reduces Acoustic Startle Response in *Bsn^Dlx5/6^* cKO Mice and Causes Prepulse Inhibition Deficits in *Bsn^Dlx5/6^* cKO and *Bsn^Emx1^* cKO Mice

To characterize baseline acoustic startle reactivity and facilitate the interpretation of potential PPI phenotypes, we first measured the startle magnitude to acoustic stimuli with varying intensities. Compared to their wildtype littermates, *Bsn^Dlx5/6^*cKO mice exhibited a significantly reduced ASR, with the difference becoming more pronounced at higher stimulus intensities (Figure 2A; ANOVA: Genotype: *F*_(1,37)_ = 4.18, *p* = 0.048; Intensity: *F*_(1.687, 62.43)_ = 51.11, *p* < 0.0001; Interaction: *F*_(7,259)_ = 2.68, *p* = 0.01). However the effect did not reach statistical significance at any of the stimulus intensities (Sidak’s multiple comparisons: 78 dB SPL: *t*_(35.65)_ = 0.25; *p* > 0.99; 84 dB SPL: *t*_(36.99)_ = 1.49, *p* = 0.71; 90 dB SPL: *t*_(36.98)_ = 1.32, *p* = 0.82; 96 dB SPL: *t*_(36.97)_ = 1.71, *p* = 0.55; 102 dB SPL: *t*_(33.16)_ = 2.73, *p* = 0.08; 108 dB SPL: *t*_(35.10)_ = 1.26, *p* = 0.86; 114 dB SPL: *t*_(28.95)_ = 1.85, *p* = 0.47; 120 dB SPL: *t*_(29.74)_ = 2.07, *p* = 0.32). The other two *Bsn* cKO strains showed no significant alterations in startle magnitude when compared to their wildtype littermates: *Bsn^Emx1^*cKO (Figure 2D; ANOVA: Genotype: *F*_(1, 53)_ = 0.0001, *p* = 0.99; Intensity: *F*_(2.192, 116.2)_ = 89.31, *p* < 0.0001; Interaction: *F*_(7, 371)_ = 0.34, *p* = 0.93); *Bsn^DAT^*cKO (Figure 2G; ANOVA: Genotype: *F*_(1,39)_ = 0.33, *p* = 0.57; Intensity: *F*_(3.231, 126.0)_ = 87.69, *p* < 0.0001; Interaction: *F*_(7, 273)_ = 0.40, *p* = 0.90).

**Figure 2:**
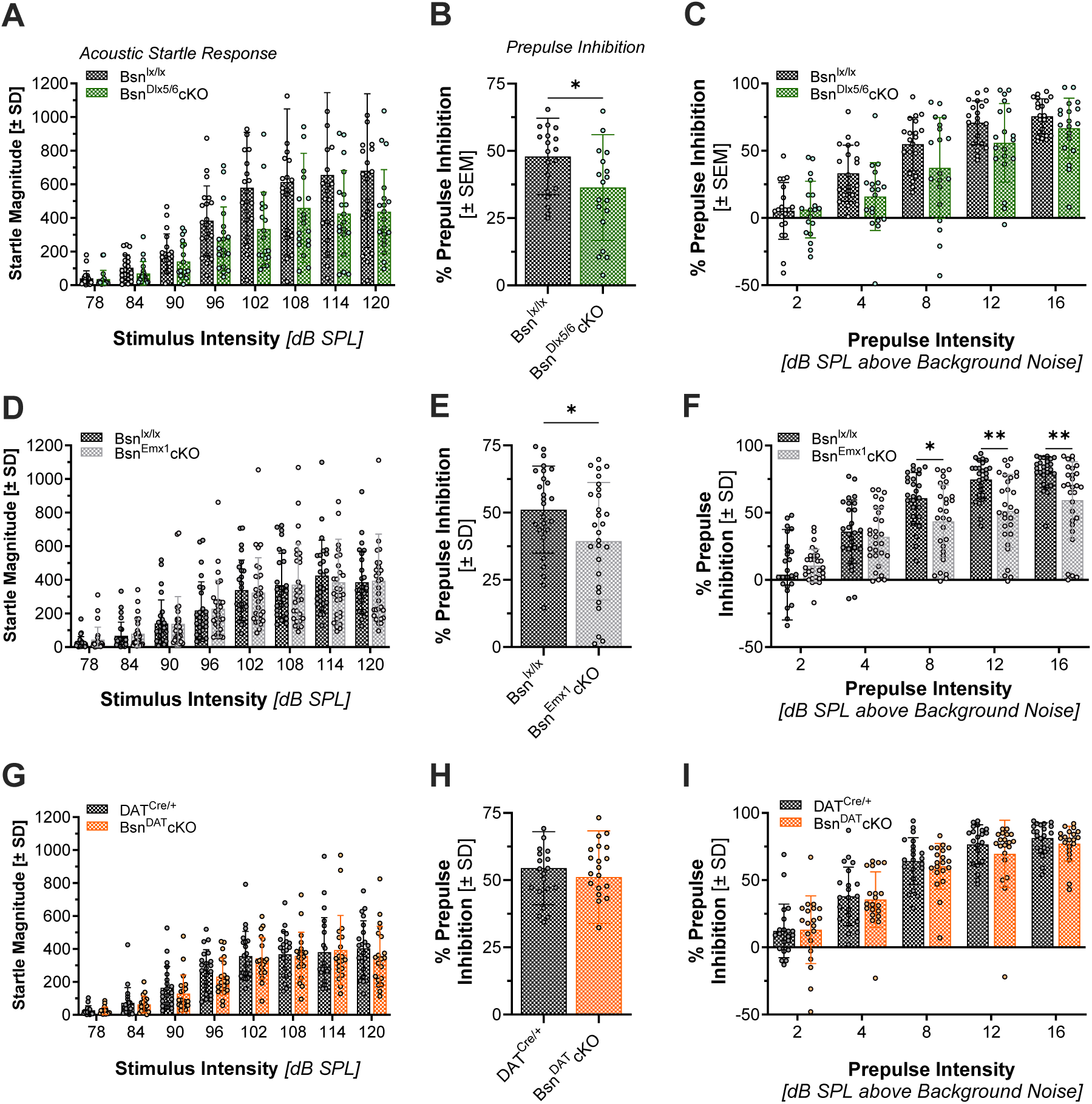
Acoustic Startle Response (ASR) and Prepulse Inhibition (PPI) in three different conditional *Bsn* knockout mice. In *Bsn^Dlx5/6^* cKO mice, *Bsn* deficiency at inhibitory synapses resulted in a significant reduction in ASR (A). This genotype also induced PPI deficits (B, C). Startle magnitude was not significantly affected by genotype in either *Bsn^Emx1^*cKO or *Bsn^DAT^* cKO mice compared to their wildtype littermates (D, G). While both DATcre/+ and *Bsn^DAT^* cKO mice exhibited similarly lowered PPI (H, I), *Bsn^Emx1^* cKO mice showed reduced PPI and differed significantly form their control littermates (E, F). The genotype effect and its interaction with prepulse intensity were most evident at higher prepulse levels (F). The bar diagrams show the mean + standard error of the mean (SEM). **p < 0.01, *p < 0.05; comparisons as indicated, following significant main effects and/or interactions in ANOVA.

PPI is widely used as an operational measure for sensorimotor gating and describes the reduction of the startle magnitude when the startle stimulus is preceded by a weak (non-startling) sensory stimulus (prepulse) (Braff, 1992).

We found overall PPI deficits in *Bsn^Dlx5/6^*cKO (Figure 2B: t-test: *t*_(37)_ = 2.11, *p =* 0.04; Figure 2C: ANOVA: Genotype: *F*_(1, 37)_ = 4.45, p = 0.04) and *Bsn^Emx1^*cKO mice (Figure 2E: t-test: *t*_(53)_ = 2.24, *p =* 0.03; Figure 2F: ANOVA: Genotype: *F*_(1,53)_ = 5.02, p = 0.03. As expected, PPI was more pronounced with higher prepulse intensities in *Bsn^Dlx5/6^*cKO mice (Figure 2C; ANOVA: Prepulse Intensity: *F*_(2.57, 95.07)_ = 89.51, p < 0.0001; Interaction: *F*_(4,148)_ = 1.86, *p* = 0.12; Genotype: *F*_(1, 37)_ = 4.45, p = 0.04).

For *Bsn^Emx1^*cKO (Figure 2F; ANOVA: Prepulse Intensity: *F*_(1.736, 91.98)_ = 176.9, *p <* 0.0001; Interaction: *F*_(4, 212)_ = 11.46, *p <* 0.0001) a similar increase was observed. Post-hoc comparisons showed significantly reduced PPI after prepulses with the intensity of 8, 12 and 16 dB SPL above background noise (Sidak’s multiple comparisons: 2 dB SPL: *t*_(30.85)_ = 0.97; *p* = 0.87; 4 dB SPL: *t*_(50.90)_ = 0.62, *p* = 0.97; 8 dB SPL: *t*_(50.30)_ = 2.74, *p* = 0.04; 12 dB SPL: *t*_(43.31)_ = 4.05, *p* = 0.001; 16 dB SPL: *t*_(37.63)_ = 3.63, *p* = 0.004).

In *Bsn^DAT^*cKO mice, no genotype or intensity effect on PPI in the cKOs were found (Figure 2H: t-test: *t*_(39)_ = 0.68, *p =* 0.50; Figure 2I: ANOVA: Genotype: *F*_(1,39)_ = 0.47, p = 0.50; Prepulse Intensity: *F*_(3.142, 122.6)_ = 201.8, p < 0.0001; Interaction: *F*_(3.142, 122.6)_ = 0.54, *p* = 0.66).

### *Bsn* Deficiency Affects Safety Learning in a Neuron Type-specific Manner but Does Not Alter Contextual Fear Expression

To investigate conditioned safety, mice were first submitted to an acclimation session in which freezing behavior to a novel context and the presentation of a novel tone stimulus (the to-be-conditioned safety CS) was evaluated. In all three strains, cKO mice, as well as their wildtype littermates froze more to the tone stimulus than to the novel context indicating weak unconditioned fear to the tone stimulus: *Bsn^Dlx5/6^*cKO mice exhibited significantly increased freezing behavior in response to the novel tone stimulus compared with control mice (Figure 3A; ANOVA: Trial type (context vs. tone): *F*_(1,37)_ = 14.61, *p* = 0.0005; Genotype: *F*_(1,37)_ = 1.39, *p* = 0.25; Interaction: *F*_(1,37)_ = 6.20, *p* = 0.01). Post-hoc Sidak’s multiple comparison analysis revealed that *Bsn^Dlx5/6^*cKO mice exhibited significantly higher freezing during the tone than during the context period (*t*_(37)_ = 4.41, *p* = 0.0002), whereas control mice showed no significant difference in freezing between the context and the tone (*t*_(37)_ = 0.95, *p* = 0.57).

**Figure 3:**
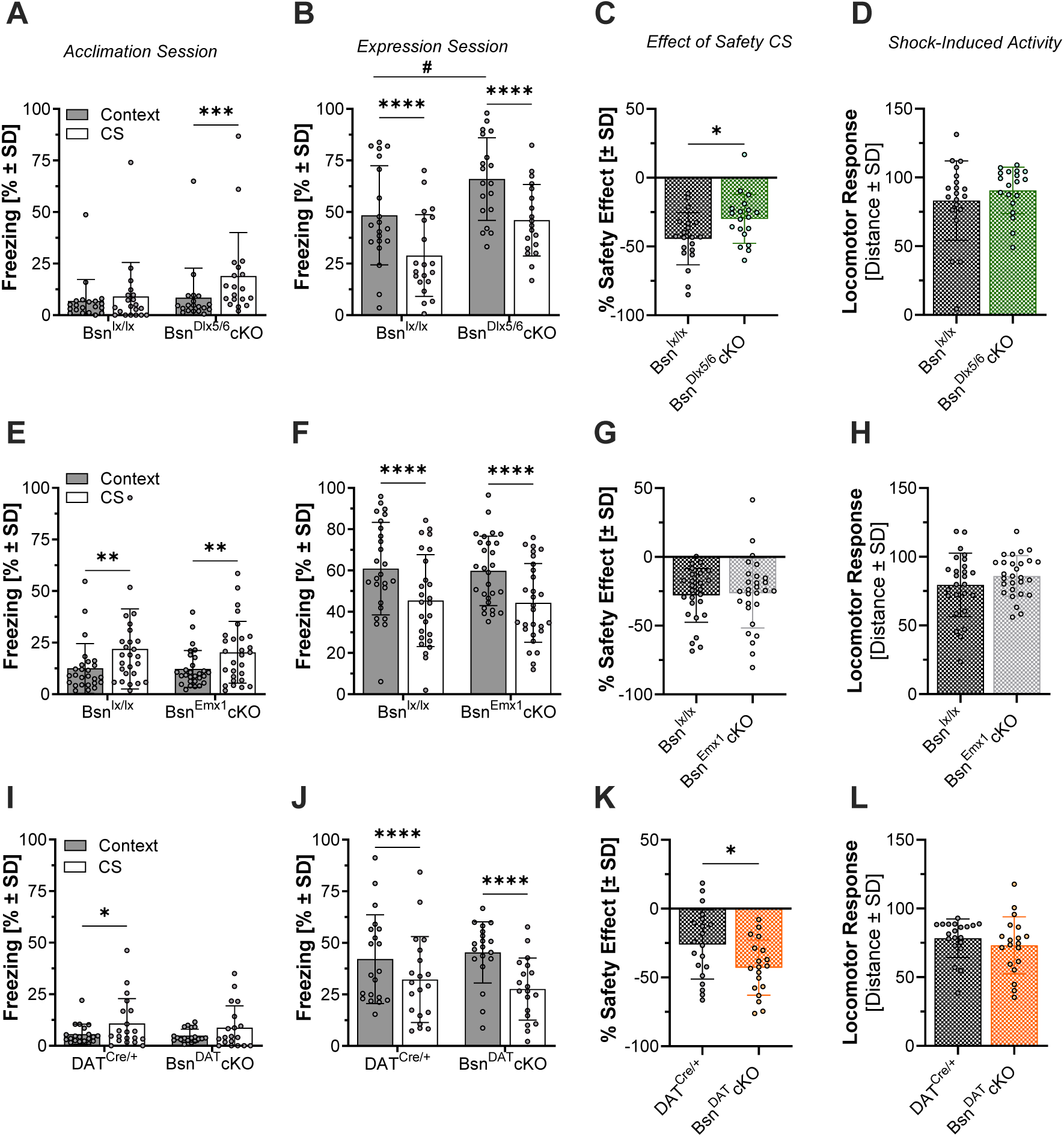
*Bsn* deficiency in *dlx5/6*-controlled GABAergic neurons decreases contextual fear and blocks conditioned safety learning while *Bsn* deficiency in dopaminergic neurons enhances safety learning. During the acclimation session, all mouse strains froze more to the tone than to the context, indicating mild unconditioned fear to the tone (A, E and I). After fear/safety conditioning, all mice successfully reduced freezing during presentations of the safety CS compared with the threat-associated context (B, F and J). However, compared to wildtype littermate controls, *Bsn^Dlx5/6^*cKO mice showed impaired safety learning (C), whereas *Bsn^Emx1^*cKO mice showed no change in safety learning (G), and *Bsn^DAT^*cKO mice showed enhanced safety learning (K). In addition, there was no genotype effect on the locomotor reactivity to the aversive foot shocks (D, H and L). Bar graphs represent mean ± standard error of the mean (SEM) of context-(black bars) and safety CS (white bars)-induced freezing behavior during an acclimation session before conditioning and a memory recall (expression) session, as well as the percent effect of the safety CS on freezing, and the locomotor response to the electric stimuli. ****p < 0.0001, ##,**p < 0.01, #,*p < 0.05; comparisons as indicated.

*Bsn^Emx1^*cKO mice also showed such an increased freezing response during the tone albeit the same was observed in its paired controls (Figure 3E; ANOVA: Trial type: *F*_(1,52)_ = 20.97, *p* < 0.0001; Genotype: *F*_(1,52)_ = 0.08, *p* = 0.78; Interaction: *F*_(1,52)_ = 0.13, *p* = 0.71; Post-hoc Sidak’s multiple comparison: *Bsn^lx/lx^*: *t*_(26)_ = 3.43, *p* = 0.002; *Bsn^Emx1^*cKO: *t*_(28)_ = 3.04, *p* = 0.008).

Both, *Bsn^DAT^*cKO and control mice exhibited increased freezing behavior in response to the tone (Figure 3I; ANOVA: Trial type: *F*_(1,37)_ = 8.29, *p* = 0.006; Genotype: *F*_(1,37)_ = 0.36, *p* = 0.55; Interaction: *F*_(1,37)_ = 0.14, *p* = 0.71); however, post-hoc analyses revealed that this increase reached statistical significance only in control mice (Post-hoc Sidak’s multiple comparison: *DAT^Cre/+^*: *t*_(20)_ = 2.33, *p* = 0.05; *Bsn^DAT^*cKO: *t*_(19)_ = 1.75, *p* = 0.17).

Following the acclimation session, mice underwent fear/safety conditioning on two consecutive days, during which the tone (safety CS) was repeatedly presented in the absence of the aversive stimulus, while the conditioning context was associated with foot shock. On the following day, learned safety was assessed in the expression session by comparing freezing during the threat-associated context with freezing during safety CS. Freezing was significantly reduced during presentation of the safety CS compared with the context in both cKO mice and wildtype littermates, across all three cKO strains:

*Bsn^Dlx5/6^*cKO mice exhibited generally increased freezing during both the threat-associated context and presentation of the safety CS compared with control mice (Figure 3B; ANOVA: Genotype: *F*_(1,37)_ = 7.63, *p* = 0.009). Nevertheless, both genotypes showed significantly decreased freezing during the safety CS compared with the threat-associated context (Trial type: *F*_(1,37)_ = 107.0, *p* < 0.0001; Interaction: *F*_(1,37)_ = 0.01, *p* = 0.90). Post-hoc Sidak’s multiple comparison confirmed the significant reduction in freezing behavior during presentations of the safety CS in both genotypes (Control: *t*_(37)_ = 7.32, *p* < 0.0001; cKO: *t*_(37)_ = 7.32, *p* < 0.0001). In addition, *Bsn^Emx1^*cKO and control mice also showed a significant decrease in freezing during presentation of the safety CS compared with the threat-associated context (Figure 3F; ANOVA: Trial type: *F*_(1,52)_ = 73.02, *p* < 0.0001; Interaction: *F*_(1,52)_ = 0.00009537, *p* = 0.99). Post-hoc Sidak’s multiple-comparisons test confirmed this reduction in both control (*Bsn^lx/lx^*: *t*_(26)_ = 5.93, *p* < 0.0001) and *Bsn^Emx1^*cKO mice (*t*_(28)_ = 6.17, *p* < 0.0001). There was no main effect of genotype on freezing behavior (Genotype: *F*_(1,52)_ = 0.04, *p* = 0.83).

Similarly, *Bsn^DAT^*cKO and DAT^Cre/+^ control mice also exhibited a significant decrease in freezing behavior during presentation of the safety stimulus compared with the threat-associated context (Figure 3J; ANOVA: Trial type: *F*_(1,37)_ = 111.4, *p* < 0.0001; Interaction: *F*_(1,37)_ = 9.08, *p* = 0.005). Also here, post-hoc Sidak’s multiple comparison test in *Bsn^DAT^*cKO mice confirmed that both cKO and control mice significantly reduced freezing in response to the safety stimulus (DAT^Cre/+^: *t*_(37)_ = 5.40, *p* < 0.0001; cKO: *t*_(37)_ = 9.47, *p* < 0.0001). There was no main effect of genotype on freezing behavior (Genotype: *F*_(1,37)_ = 0.01, *p* = 0.90).

Due to the substantial inter-individual variability and genotype-dependent differences in overall freezing levels, percent difference scores were calculated for each individual to quantify the magnitude of safety learning relative to their contextual fear response. These percent differences scores confirmed that there was reduced freezing behavior during the safety CS in all groups. However, *Bsn^Dlx5/6^*cKO exhibited a significantly reduced safety learning effect compared to their littermate controls (Figure 3C; t-test: *t*_(37)_ = 2.46, *p =* 0.01). In contrast, no significant difference was observed between *Bsn^Emx1^*cKO and control littermates (Figure 3G; t-test: *t*_(52)_ = 0.25, *p =* 0.80), while *Bsn^DAT^*cKO mice showed an enhanced safety learning effect compared with control mice (Figure 3K; t-test: *t*_(37)_ = 2.30, *p =* 0.03).

Since fear and safety learning can also be modulated by the reactivity to the used aversive stimuli, we also analyzed the locomotor response of the mice to the electric stimuli. This locomotor response was not affected by the genotype across all three strains (*Bsn^Dlx5/6^*cKO: Figure 3D: t-test: t_(37)_ = 0.97, p = 0.34; *Bsn^Emx1^*cKO: Figure 3H: t-test: t_(52)_ = 1.26, p = 0.22; *Bsn^DAT^*cKO: Figure 3L: t-test: t_(37)_ = 0.91, p = 0.37), indicating that the observed effects on safety learning were not due to enhanced or reduced processing of the aversive stimuli.

### Elevated Anxiety-like Behavior in *Bsn*-Deficient Mice in the Light/Dark Box Test

The light/dark box test is a widely used behavioral assay to assess anxiety-like behavior in rodents, based on their natural aversion to brightly illuminated areas (Bourin and Hascoët, 2003).

Compared to their wildtype littermates, *Bsn^Dlx5/6^* cKO mice exhibited increased anxiety-like behaviors, characterized by significant reduced time spent in the light compartment (Figure 4A; t-test: *t*_(37)_ = 6.38, *p <* 0.0001), reduced distance travelled in the light compartment (Figure 4B; t-test: *t*_(37)_ = 6.49, *p <* 0.0001), and fewer entries into the light compartment (Figure 4C; t-test: *t*_(37)_ = 2.65, *p =* 0.01). Moreover, *Bsn^Dlx5/6^* cKO mice displayed increased overall locomotor activity, as indicated by greater total distance travelled (Figure 4D; t-test: *t*_(37)_ = 3.23, *p =* 0.003).

**Figure 4:**
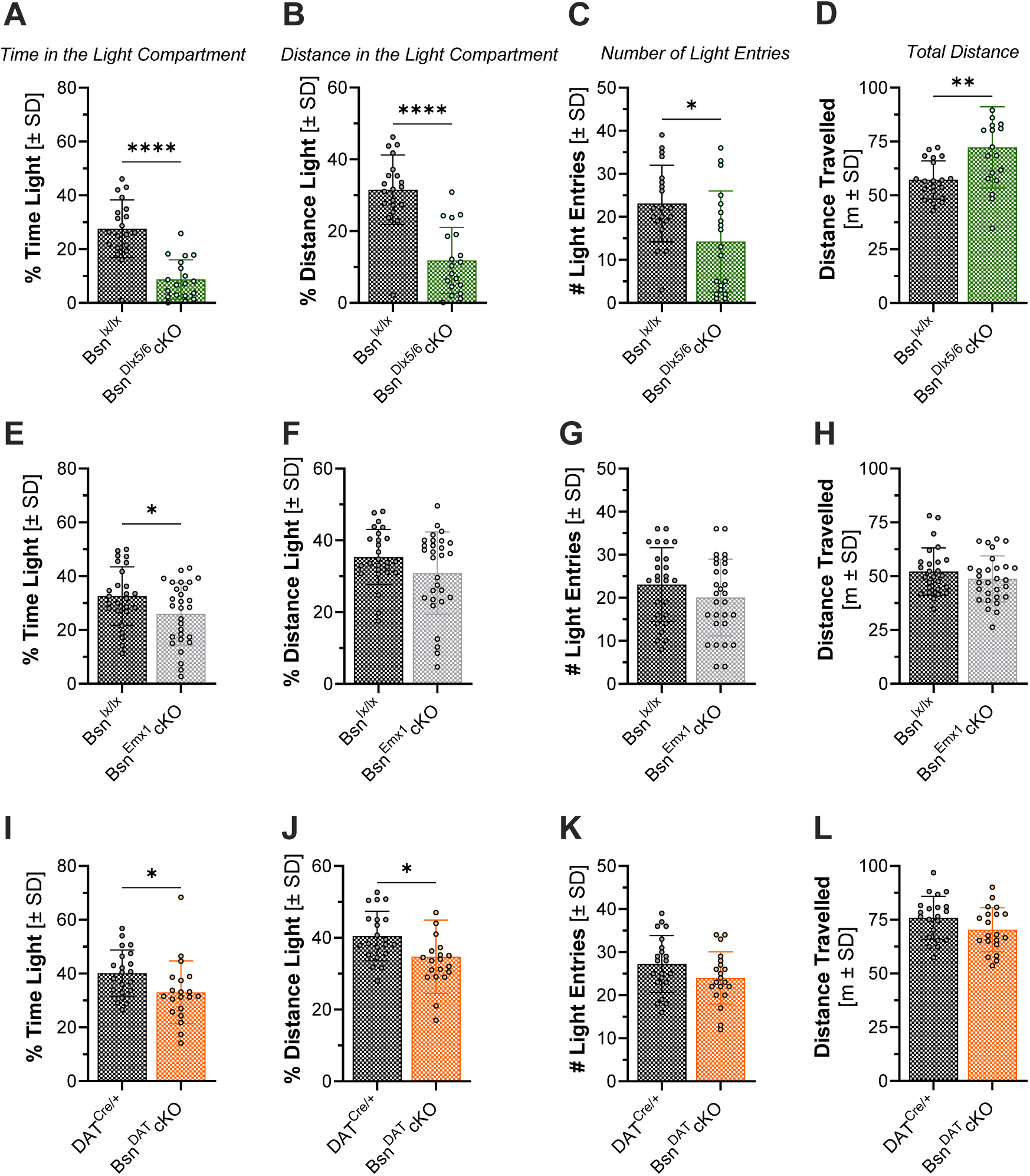
*Bsn* deficiency increases anxiety-like behavior in the light/dark box test. Compared with wildtype littermates, *Bsn^Dlx5/6^*cKO mice exhibited significantly reduced percent time spent in the light compartment (A), decreased distance traveled within the light compartment (B), and fewer entries into the light compartment (C), despite an overall increase in total distance travelled (D). *Bsn^Emx1^*cKO mice showed a significant reduction in time spent in the light compartment (E), while reductions in other measures did not reach statistical significance (F, G, H). In *Bsn^DAT^*cKO mice, both the percentage of time (I) and distance spent in the light compartment were significantly decreased (J), while total distance travelled and number of light entries showed no differences (K, L). Bar graphs represent mean ± standard error of the mean (SEM). ****p < 0.0001, **p < 0.01, *p < 0.05; comparisons as indicated.

*Bsn^Emx1^*cKO mice spent significantly less time in the light compartment compared with control mice (Figure 4E; t-test: *t*_(53)_ = 2.17, *p =* 0.03), while no significant genotype differences were observed for the distance travelled in light compartment (Figure 4F: t-test: *t*_(53)_ = 1.68, *p =* 0.099),the number of entries into the light compartment (Figure 4G: t-test: *t*_(53)_ = 1.27, *p =* 0.21) or the total distance travelled (Figure 4H: t-test: *t*_(53)_ = 1.15, *p =* 0.26).

In *Bsn^DAT^* cKO mice, both the percentage of time spent in the light compartment (Figure 4I; t-test: *t*_(39)_ = 2.22, *p =* 0.03) and distance (Figure 4J; t-test: *t*_(39)_ = 2.12, *p =* 0.04) spent in the light compartment were significantly decreased compared with their control mice. The number of entries into the light compartment (Figure 4K; t-test: *t*_(39)_ = 1.63, *p =* 0.11) and the total distance travelled (Figure 4L; t-test: *t*_(39)_ = 1.76, *p =* 0.09) did not differ significantly from wildtype controls.

### Absence of *Bsn* Results in Mild to Severe Alterations in Brain Volume in a Neuron Type-Specific Manner

Global absence of Bassoon in null mutants resulted in a surprising volume increase mainly in forebrain regions that is steadily ongoing during brain maturation and not observed for the cerebellum (Angenstein et al., 2007; Heyden et al., 2011). To assess which type of neurons might contribute to this expansion, we performed volume measurements utilizing 9.4T-MRI. The absence of Bassoon in different subtypes of synapses consistently led to an increase in volume in the hippocampus, albeit to varying degrees.

*Bsn^Dlx5/6^*cKO mice showed a substantial increase in total brain volume compared to control mice (Figure 5A). These mice exhibited significant increases in cortical volume (Figure 5B; t-test: t_(10)_ = -4.47, p < 0.001), hippocampal volume (∼25% increase, Figure 5C; t-test: t_(10)_ = -5.11, p < 0.0001) and striatal volume (Figure 5D; t-test: t_(10)_ = -3.59, p < 0.005).

**Figure 5:**
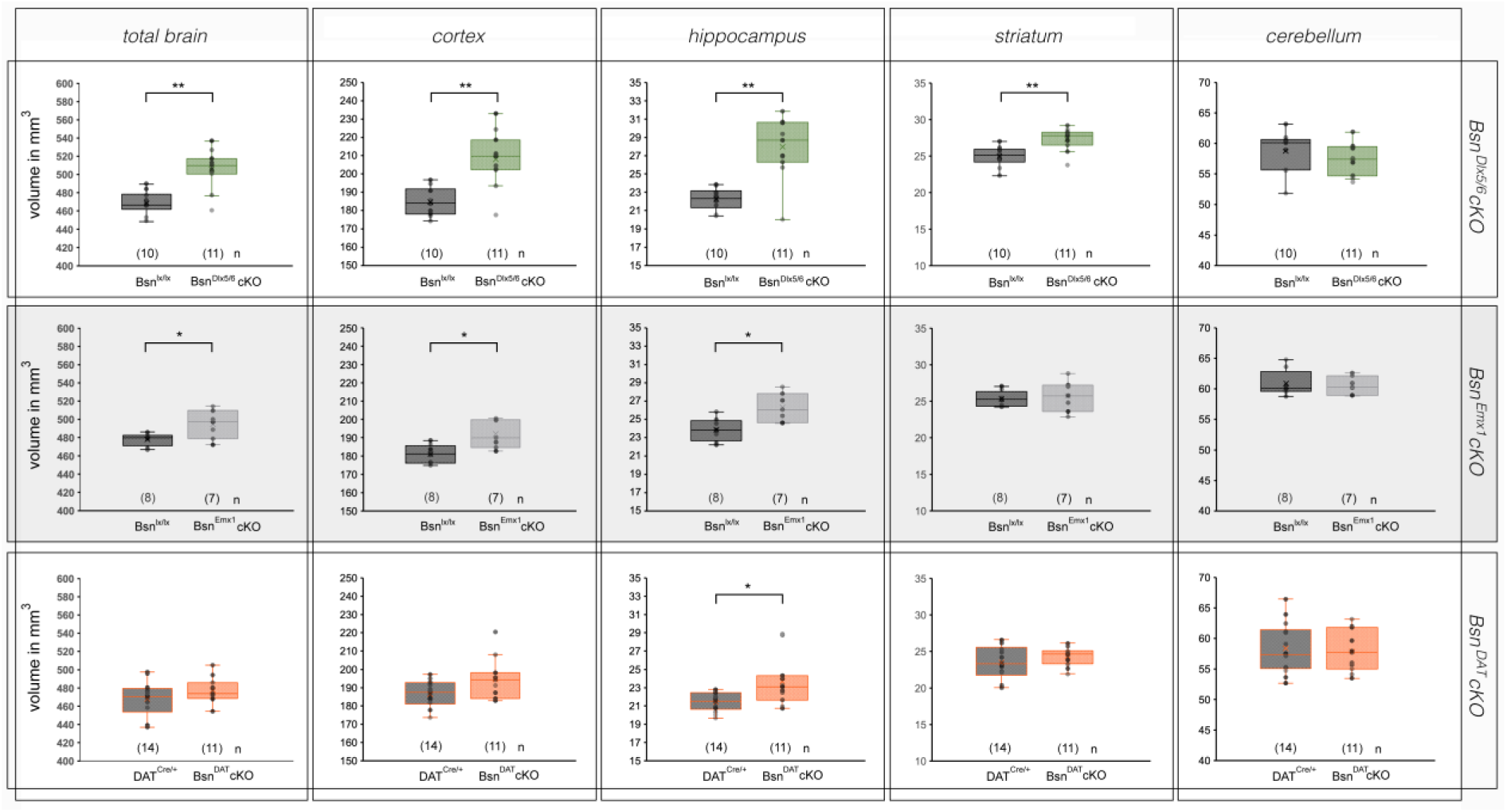
Anatomical changes in *Bsn* cKO mice: Summary of volumetric changes in selected brain regions of mice that lack *Bsn* in distinct inhibitory (*Bsn^Dlx5/6^*cKO, A-E), excitatory (*Bsn^Emx1^*cKO, F-J) or dopaminergic neurons (*Bsn^DAT^*cKO, K-O). Significant differences are indicated an asterisks (* p < 0.01, ** p < 0.001).

By contrast, *Bsn^Emx1^*cKO mice showed a moderate increase in the total brain volume (Figure 5F). Significant increases were observed in cortical volume (Figure 5G; t-test: t_(7)_ = -3.36, p < 0.01) and hippocampal volume (Figure 5H; t_(7)_ = -3.36, p < 0.01), but not the in striatal volume (Figure 5I).

*Bsn^DAT^*cKO mice showed no significant changes on total brain volume (Figure 5K) or on cortical volume (Figure 5 L; t-test: t_(10)_ = -1.97, p = 0.061), but exhibited a significant increase in hippocampal volume (Figure 5M; t-test: t_(10)_ = -2.84, p < 0.01). Striatal (Figure 5N) and cerebellar volume remained unaffected. Notably, cerebellar volume was comparable between each of the three *Bsn* cKO genotypes and their corresponding controls (Figure 5 E, J, O).

## DISCUSSION

Symptoms of schizophrenia include positive (hallucinations, delusions), negative (social withdrawal, flat affect and lack of motivation) aspects and cognitive disturbances, which has been associated with dysregulation of dopaminergic, glutamatergic, and GABAergic signaling (Howes et al., 2017; Jahangir et al., 2021; Kruse and Bustillo, 2022; McCutcheon et al., 2023). The aim of the present study was to dissect the contribution of different neurotransmitter systems to the development of schizophrenia-related behavioral phenotypes elicited by mutation of the *Bsn* gene. We focused on three neuronal populations: (1) GABAergic interneurons, whose development depends on expression of the *dlx5/6* gene enhancer (*Bsn^Dlx5/6^*cKO), (2) glutamatergic forebrain neurons (*Bsn^Emx1^*cKO), and (3) dopaminergic neurons (*Bsn^DAT^*cKO). Specifically, we assessed whether these manipulations affect behavioral domains commonly disrupted in individuals with schizophrenia and frequently modeled in mouse studies of the disorder. We found that both *Bsn^Dlx5/6^*cKO and *Bsn^Emx1^*cKO mice exhibited deficits in sensorimotor gating, as indicated by reduced PPI, whereas *Bsn^DAT^*cKO showed no alterations. Notably, PPI is calculated relative to the individual startle response, thereby accounting for differences in baseline startle magnitude. Although *Bsn^Dlx5/6^*cKO mice exhibited a reduced startle magnitude, the concomitant reduction in PPI indicates impaired sensorimotor gating that cannot be explained solely by their lower baseline startle response. In the fear/safety discrimination paradigm, *Bsn^Dlx5/6^*cKO mice exhibited impaired safety learning, whereas *Bsn^DAT^*cKO mice showed enhanced safety discrimination. All three cKO lines displayed increased anxiety-like behavior in light/dark box test. Finally, both the *Bsn^Dlx5/6^*cKO and *Bsn^Emx1^*cKO mice showed increased total brain volume, associated with enlargement of specific brain regions, including the cortex and hippocampus, similar to the volumetric alterations reported in constitutive *Bsn* mutants (Angenstein et al., 2007; Heyden et al., 2011).

Synaptic dysfunction has long been recognized as a core feature of schizophrenia, with evidence from pharmacological studies implicating dopaminergic, glutamatergic, and GABAergic neurotransmission, including dopamine D2, NMDA, and GABA_A_ receptors. Advances in genome-wide association studies have further highlighted the genetic underpinnings of the disorder, identifying numerous synaptic genes with single nucleotide polymorphisms and other variants associated with schizophrenia. These findings have led to the proposal that schizophrenia can be conceptualized as a synaptopathy (Kenny et al., 2014; Pocklington et al., 2014). Among the synaptic proteins implicated in the disorder, Bassoon has emerged as a molecule of particular interest (Chen et al., 2021; Martins-de-Souza et al., 2009; Popiolek et al., 2011).

In our study, *Bsn^DAT^*cKO mice demonstrated enhanced safety learning, whereas *Bsn^Dlx5/6^*cKO mice displayed impaired safety learning. These effects are unlikely to be attributable to differences in sensory perception of the safety signal, contextual cues, or sensitivity to the aversive stimulus itself, as all three cKO lines exhibited normal responses to the safety cue during acclimation, intact contextual fear conditioning, and comparable locomotor responses to the aversive foot shocks. The opposing effects observed in fear/safety learning across the *Bsn* cKO lines likely reflect circuit-specific consequences of Bassoon loss. In *Bsn^Dlx5/6^*cKO mice, deletion of Bassoon from GABAergic neurons may weaken inhibitory control within cortical and limbic networks. Such alteration in excitation-inhibition balance could compromise the temporal coordination and synchronization of neural activity required for appropriate suppression of conditioned fear during safety cue presentation, thereby contributing to reduced safety learning. In contrast, the mechanisms underlying the enhanced safety learning in *Bsn^DAT^*cKO mice remain unclear. Given the established involvement of dopaminergic signaling in threat and safety learning, altered dopamine-dependent modulation of associative learning could contribute to this phenotype. However, whether Bassoon deletion in dopaminergic neurons alters dopamine release or signaling remains to be determined. Together, these bidirectional behavioral effects emphasize the importance of Bassoon in maintaining the functional integrity of distinct neural circuits involved in fear and safety learning.

The complex interplay between dopaminergic and glutamatergic systems, including their co-release and the modulatory role of GABA in maintaining the dopamine-glutamate balance within the cortico-striato-thalamo-cortical loop, has been implicated in the pathophysiology of schizophrenia (Buck et al., 2022; McCutcheon et al., 2023). Sensorimotor gating, i.e. the ability to filter or inhibit responses to irrelevant sensory information, can be assessed using PPI of the ASR, which is consistently impaired in individuals with schizophrenia and in corresponding animal models. Accordingly, PPI is widely used as a translational measure of schizophrenia-related sensorimotor-gating deficits (Mena et al., 2016; Sato, 2020). In the present study, *Bsn^Dlx5/6^* cKO and *Bsn^Emx1^* cKO mice displayed impaired PPI, whereas *Bsn^DAT^* cKO mice did not show any alterations relative to their respective controls. However, interpretation of the PPI phenotype requires consideration of baseline startle reactivity, as PPI is calculated relative to the individual startle response. In particular, *Bsn^Dlx5/6^*cKO mice exhibited reduced ASR magnitude, which may complicate the interpretation of their reduced PPI. Thus, the observed PPI phenotype should be interpreted in the context of altered baseline startle reactivity rather than as evidence of an isolated deficit in sensorimotor gating. PPI is sensitive to pharmacological manipulation of glutamatergic, GABAergic or dopaminergic neurotransmission. In particular, antagonism of glutamatergic NMDA or GABA_B_ receptors, as well as activation of dopamine receptors through direct or indirect agonists, has been shown to disrupt PPI (Linderholm et al., 2010; Yeomans et al., 2010; Zhang et al., 2000). Thus, the PPI deficits observed in *Bsn^Emx1^* cKO and *Bsn^Dlx5/6^* cKO mice is therefore consistent with the involvement of excitatory-inhibitory and neuromodulatory circuits in the regulation of sensorimotor gating, although the underlying mechanisms remain to be determined.

A deficit in safety learning has been reported in individuals with schizophrenia (Holt et al., 2012), potentially reflecting dysfunction of the prefrontal cortex – a brain region critically involved in the expression of conditioned safety memory (Kreutzmann et al., 2020a; Kreutzmann and Fendt, 2020). Of note, schizophrenia has been associated with disrupted dopaminergic (Knable and Weinberger, 1997) and GABAergic signaling (Lewis, 2000) within the prefrontal cortex, while findings on glutamatergic transmission in this region remain complex and, at times, inconsistent (Schoonover et al., 2020). Pharmacological studies further support the contribution of these systems: reduced dopamine signaling in the prefrontal cortex impairs safety learning (Duvarci, 2024; Yan et al., 2019), whereas decreased GABAergic signaling in the prefrontal cortex or the amygdala has been shown to facilitate it (El Matine et al., 2023; Kreutzmann and Fendt, 2020). These observations appear, at first glance, to contrast with our results, in which *Bsn* deletion in GABAergic or dopaminergic neurons produced opposite outcomes. However, dopaminergic dysfunction has been proposed to confer aberrant salience to otherwise non-threatening or neutral cues (Kapur, 2003), reflecting altered prediction error signaling rather than changes in learning capacity *per se* (Corlett et al., 2016, 2007). Consistent with this view, psychosis-prone and paranoid phenotypes show exaggerated behavioral and neural responses to non-threatening cues, including safety signals, suggesting that increased responding may reflect heightened cue salience or predictive value rather than improved safety discrimination (Jensen et al., 2008; Powers et al., 2017). Within this framework, *Bsn* deletion in dopaminergic neurons may enhance the perceived salience or predictive value of safety cues, thereby increasing behavioral responding despite intact - or even impaired - discriminative learning.

Anxiety-related symptoms are highly prevalent in schizophrenia, affecting approximately 65% of patients, with social anxiety disorder being the most frequent comorbid condition (Temmingh and Stein, 2015). In the light/dark box test, all three *Bsn* cKO lines exhibited enhanced anxiety-like behavior compared with their respective controls, with the most pronounced phenotype observed in *Bsn^Dlx5/6^*cKO mice. This finding is consistent with previous results from our group showing increased anxiety-like behavior in male *Bsn^Dlx5/6^*cKO mice under different light conditions during the active phase of the light/dark cycle (Annamneedi et al., 2026). The replication of this phenotype across different light conditions further supports a robust anxiety-related phenotype in *Bsn^Dlx5/6^*cKO mice. Dysregulation of the GABAergic system, particularly involving parvalbumin-positive interneurons across cortical and subcortical regions, has been implicated in the emergence of anxiety-like symptoms, as well as in the broader emotional and cognitive disturbances observed in schizophrenia and corresponding mouse models (Gawande et al., 2023; Taylor et al., 2014), including *Bsn^Dlx5/6^*cKO mice (Annamneedi et al., 2026). Moreover, reductions in glutamatergic or dopaminergic signaling, either induced pharmacologically or resulting from conditions such as Parkinsons’s disease or schizophrenia, have also been shown to elicit anxiety-like states in rodents (De La Mora et al., 2010; Durant et al., 2009; Plaznik et al., 1994).

Finally, we conducted a comparative analysis of brain volumes in the three *Bsn* cKO mouse strains using MRI. Both *Bsn^Dlx5/6^*cKO and *Bsn^Emx1^*cKO mice exhibited increased total brain volume, with significant enlargements in the cortex and hippocampus. In contrast, the *Bsn^DAT^*cKO mice showed a more selective effect, with increased volume observed specifically in the hippocampus. The absence of functional *Bsn* in all neurons has previously been associated with a substantial enlargement of the hippocampus and cortex (Angenstein et al., 2007; Heyden et al., 2011). On the other hand, human studies have consistently linked schizophrenia to volume reductions in the hippocampus but enlarged ventricles (Haukvik et al., 2015; Reif et al., 2006; Svancer and Spaniel, 2021) and a study reported increased head circumference (Bassett et al., 1996). This volumetric increase observed in *Bsn* cKO mice particularly evident in hippocampus, might be attributed to elevated brain-derived neurotrophic factor (BDNF) levels in the hippocampus and other forebrain structures (Annamneedi et al., 2021; Dieni et al., 2012; Heyden et al., 2011). The current findings suggest that the hippocampal volume increase is primarily driven by the absence of Bassoon in inhibitory synapses, whereas loss of Bassoon in excitatory or dopaminergic synapses appears to have less impact. Notably, both the *Bsn^Dlx5/6^*cKO and *Bsn^Emx1^*cKO mice exhibit epilepsy, though with varying severity (Blondiaux et al., 2023) and epilepsy is a common comorbidity in schizophrenia (Adachi and Ito, 2022).

Recently identified rare pathogenic mutation in the *BSN* gene (p.Arg1087Gln) in patients with schizophrenia (Chen et al., 2021), is predicted to be deleterious and disease causing. The corresponding amino acid in rat Bassoon lies upstream of the binding site for cytoskeletal motor proteins, i.e. dynein light chains (Dlc-1, Dlc-2; aa 1360-1692). This interaction between Bassoon and Dlc proteins is essential for the transport of Piccolo-Bassoon transport vesicles (PTVs) to newly forming synapses (reviewed in (Gundelfinger et al., 2016). Previous studies revealed elevated Bassoon expression in the prefrontal cortex of post-mortem brains of schizophrenic patients along with other cytoskeletal and calcium-sensing proteins (Martins-de-Souza et al., 2009). Collectively, these findings support a critical role for Bassoon in the synaptic abnormalities underlying schizophrenia and highlight the need for further investigation into its functional implication across different neural circuits.

### Conclusion

In conclusion, this study provides evidence that the loss of the presynaptic protein Bassoon in distinct neuronal populations leads to behavioral and structural brain abnormalities relevant to schizophrenia. The divergent phenotypes observed in *Bsn^Dlx5/6^*cKO, *Bsn^Emx1^*cKO, and *Bsn^DAT^*cKO mice highlight the complexity of phenotypes associated with schizophrenia. A recent study reported sex-dependent alterations in alcohol drinking-behavior and affective behaviors in *Bsn^Emx1^*cKO and *Bsn^DAT^*cKO mice (Kalinichenko et al., 2026), while another recent study provides evidence for a role of Bassoon in memory stability and cognitive flexibility. Specifically, deletion of *Bsn* in telencephalic excitatory neurons (*Bsn^NEX^*cKO) enhanced the retention of social and non-social memories and conditioned fear memories. These mice also showed impaired extinction of fear, suggesting a reduced ability to update or suppress previously learned memories in response to new learning (Zoicas et al., 2026). Together, the cell type-specific Bassoon knockout mice provide a valuable tool for dissecting the diverse behavioral, cellular, and molecular phenotypes levels associated with Bassoon dysfunction. These findings further support a potential role of Bassoon in the etiology of schizophrenia and related neurodevelopmental disorders and highlight the broader contribution of synaptic dysfunction to the pathophysiology of neuropsychiatric disease.

## Author Contributions: CRediT

**Judith C. Kreutzmann:** Conceptualization, Formal analysis, Investigation, Writing – Original draft, review & editing, visualization. **Frank Angenstein:** Conceptualization, Formal analysis, Investigation, Writing – Original draft, review & editing, Visualization. **Anna Fejtova:** Conceptualization, Writing – Review & editing, Funding acquisition. **Oliver Stork:** Conceptualization, Formal analysis, Writing – Review & editing, Supervision, Funding acquisition. **Eckart D. Gundelfinger:** Conceptualization, Writing – Original draft, review & editing, Supervision, Funding acquisition. **Markus Fendt:** Conceptualization, Writing – Original draft, review & editing, Supervision, Funding acquisition. **Anil Annamneedi:** Conceptualization, Formal analysis, Investigation, Writing – Original draft, review & editing, visualization, Funding acquisition.

## Acknowledgement

We are very grateful to animal facility technical staff from LIN and IPT for their steady support and Dr. Carolina Montenegro-Venegas for helpful discussions and support.

