## Supplementary information for "Dissection of neuropsychiatric-relevant behavioral and structural phenotypes utilizing neuron type-specific ablation of the *Bassoon* gene in mice"

**Supplementary methods:**

Male *Bsn*$^{DAT}$cKO and *DAT*$^{Cre/+}$ (control) mice were subjected to series of behavioral experiments to assess changes in their general behavioral at the IBIO, OvGU, Magdeburg.

**Home cage activity monitoring**

Mice were monitored for 3-4 consecutive days in their home cages as described in Annamneedi et al., 2018. Activity was measured using infrared-thermo sensors (Home Cage Activity System, Coulbourn Instruments, Allentown PA), mounted on the top of each cage, which detect the changes in body heat for movement of the mice. Small movement lower limit recorded for 100 milliseconds and upper limit recorded for 500 milliseconds. Movement measured between lower and upper limits and the raw values for each 15 seconds were summed up to detect activity periods in 5 minute bins per hour (total 12 active periods per hour). The percentages of active periods were calculated per hour and the mean values for each hour per genotype over 3-4 consecutive days were plotted.

**Open field exploration**

Open field exploration was performed to assess the changes in activity and anxiety under novel environment exploration. The mice were tested in an open field for two consecutive days, in an arena measuring 50 cm (l) x 50 cm (w) x 35 cm (d), 20 minutes each. During day 1, red light (5 Lux low light conditions) was used, and a bright light (100 Lux) was used during day 2. Center (25 cm x 25 cm) region of the open field chamber was marked to analyse the anxiety-like behavior. Exploration of mice was monitored using a video-tracking system (ANY-maze, Stoelting Co, USA) interfaced with computer. Distance moved by mice and percentage of time spent in centre region were measured.

**Inverted grip strength test**

The inverted grip strength test was done to assess the muscular strength based on the ability of mice to remain hanging to the inverted cage lid. Each mouse was placed on the large rat cage lid and inverted. This inverted lid was placed at 35-40 cm height from the ground level. Bedding material or big rat cage was placed under the lid in order to protect the mouse from injuries, if fallen. Hanging time was measured for 2 minutes and expressed as % of time hanging.

**Rotarod**

Motor coordination and balance was measured in mice using rotarod system (Ugo Basile Rota-Rod model#57604). Mice were placed on a rotating drum (3 cm per single mouse; 5 mice can be tested each time) with a constant speed of 4 rpm for 2 minutes during two training sessions with 24 hours interval in between. During the test, mice were placed on accelerating rotarod from 4 to 40 rpm for 5 minutes. This was done in two sessions and average values from these two sessions were considered for measuring the duration the mice spent on accelerating rotarod and maximum speed they withstand.

**Sucrose preference test**

To test depression-like behavior, mice were tested in sucrose preference test. Protocol was adopted from Strekalova et al., 2004. Three days before the start of the experiment, mice were habituated to the 15 ml falcon tubes containing tap water. During test, for 4 days, mice were provided with two falcon tubes one containing 1% sucrose solution and other containing tap water. Care was taken to prevent probable effects of side preference; the position of two falcon tubes was switched every 24 hours. Sucrose (%) preference was calculated by measuring the amount of sucrose intake with that of total amount of sucrose and water intake.

**Fear conditioning**

Mice were tested in an auditory cued fear conditioning paradigm described previously (Annamneedi et al., 2018). The fear conditioning apparatus [16 cm (l) × 32 cm (w) × 20 cm (d)] (TSE System, Bad Homburg, Germany) comprises an acrylic glass arena with a grid floor to deliver the electric foot shock. A continuous background noise of 70 dB was maintained in a sound-proof cubicle, where entire arena was placed and connected to a computer to measure the different parameters, using photo beam system. During habituation, mice were placed in training apparatus and presented with 2 sets of six neutral acoustic stimulus (CS− 2.5 kHz, 10 s with 20 s inter stimulus intervals, ISIs) separated by a 2 min pause. 24 hours later, mice were trained by presenting 2 sets (separated by a 2 min pause) of three conditional acoustic stimulus

(CS+ 10 kHz, 10 s with 20 s ISIs) each CS+ stimulus terminating with a 1 s unconditional stimulus (US, scrambled foot shock of 0.4 mA). Contextual and cue fear memory was assessed by placing the mice in training and neutral context (new standard cage with bedding), respectively. Fear memory levels were analyzed by monitoring the freezing behavior (lack of movements except for respiration) and expressed as percentage of time spent freezing.

**Supplementary Results:**

***Bsn$^{DAT}$*cKO mice display no major behavioral alterations in comparison to the controls:**

In home cage activity monitoring, *Bsn$^{DAT}$*cKO and *DAT$^{Cre/+}$* mice display increment in their activity during dark phase, with a significant of effect of phase or day time (Supplementary Figure 1A; $F(23,299)=23.43$, $p<0.0001$, two-way repeated measures ANOVA). Interestingly, there was a significant effect of genotype ($F(1,13)=5.00$, $p=0.0435$) and genotype x phase interaction ($F(23,299)=2.10$, $p=0.0027$) observed. Phase-wise comparison of activity (%) revealed no change in activity during the light phase of the cycle (Supplementary Figure 1B; *DAT$^{Cre/+}$*: 17.46 ± 3.680%; *Bsn$^{DAT}$*cKO: 21.73 ± 2.079%; $t(13)=1.045$, $p=0.3150$), while an increased activity of *Bsn$^{DAT}$*cKO mice was observed during dark phase of the cycle in comparison to control mice (Supplementary Figure 1C; *DAT$^{Cre/+}$*: 35.22 ± 5.872%, N=7; *Bsn$^{DAT}$*cKO: 55.99 ± 5.563%, N=8; $t(13)=2.566$, $p=0.0235$, Student's t-test).

Next, *Bsn$^{DAT}$*cKO and *DAT$^{Cre/+}$* mice were tested in an open field arena to analyze novelty induced exploratory behavior and anxiety-like behavior. A significant effect of test session was observed (Supplementary Figure 1D; $F(1,13)=10.82$, $p=0.0059$, two-way repeated measures ANOVA). However, no effect of genotype ($F(1,13)=0.39$, $p=0.5445$) or genotype x test session interaction ($F(1,13)=1.58$, $p=0.2314$) was observed.

Further, the time spent in center region of the open field was compared between *Bsn$^{DAT}$*cKO and *DAT$^{Cre/+}$* mice as an indication for anxiety (Supplementary Figure 1E). No effect of genotype ($F(1,13)=1.72$, $p=0.2126$, two-way repeated measures ANOVA) or genotype x test session interaction ($F(1,13)=0.58$, $p=0.4615$) could be observed, indicating no anxiety-like behavior in *Bsn$^{DAT}$*cKO.

Depression-like behavior was analysed by employing the sucrose consumption test. Both *Bsn$^{DAT}$*cKO and *DAT$^{Cre/+}$* mice consume comparable amounts of sucrose solution (normalized to the water consumption) over the course of the test (Supplementary Figure 1F; $t(13)=1.655$, $p=0.1218$, Student's t-test).

Furthermore, motor coordination and strength was analysed using inverted grip strength and rotarod test. Both groups displayed similar time periods hanging the inverted cage (Supplementary Figure 1G; $t(13)=0.7055$, $p=0.4929$, Student's t-test).

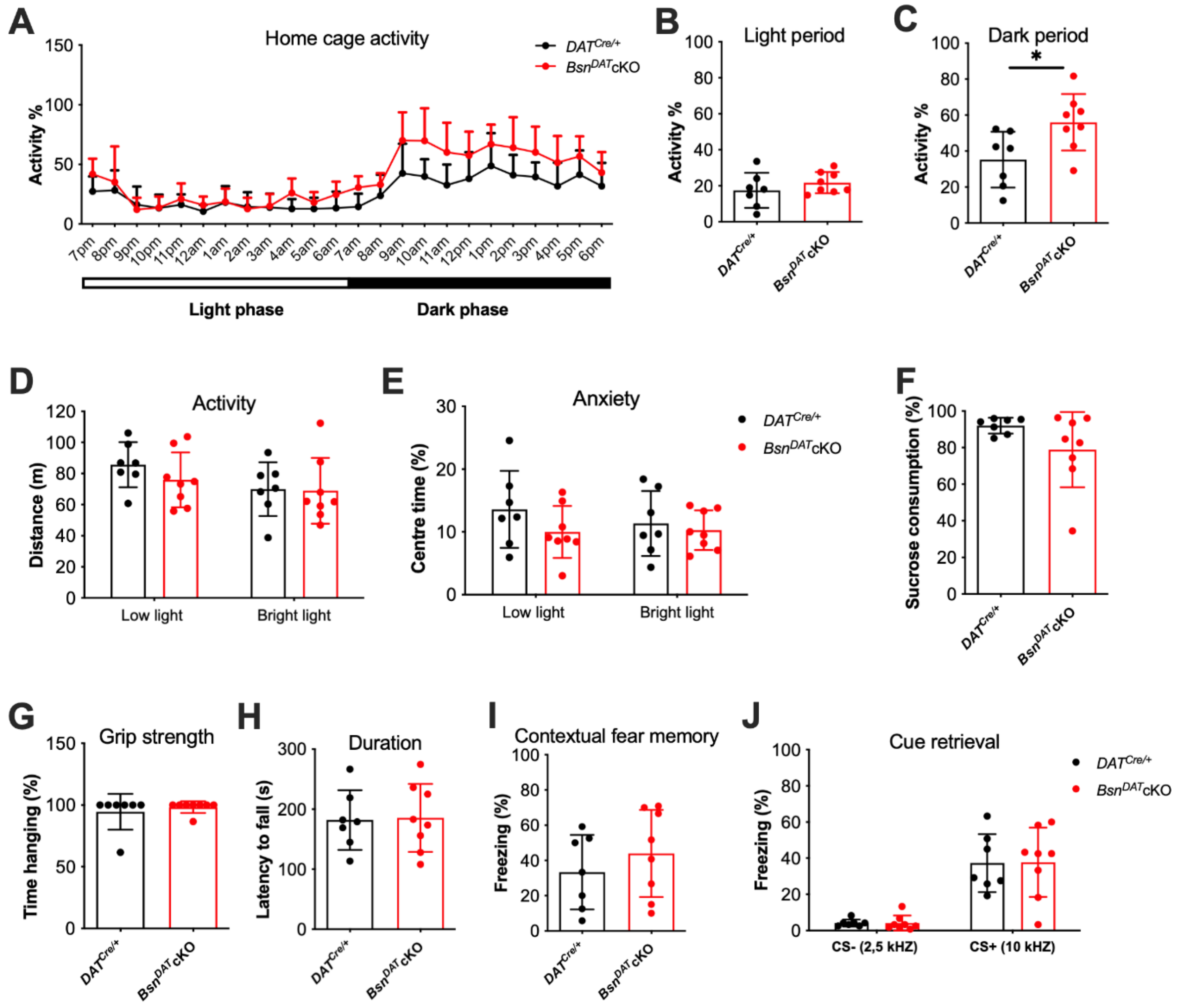


**Supplementary figure 1: Behavioral characterization of *Bsn*$^{DAT}$cKO mice.** (A-C). Home cage activity monitoring in mice, indicating normal circadian activity pattern. However, *Bsn*$^{DAT}$cKO mice display increased activity during the dark phase of the cycle. (D, E). *Bsn*$^{DAT}$cKO mice display unaltered novelty induced activity or anxiety tested in open field arena. (F). Both the groups display similar sucrose consumption rate indicating no depressive-like behavior. Normal grip strength (G) and motor functions (H) in *Bsn*$^{DAT}$cKO mice. (I,J) Classical cue-combined fear conditioning test reveals no changes in memory between the groups during contextual and cue retrieval sessions. *Bsn*$^{DAT}$cKO: N=8 and *DAT*$^{Cre/+}$: N=7; Data represented as mean ± SEM.

Next, mice were tested for motor coordination and learning using an accelerating rotarod from 4-40 rpm for 5 minutes. Both *Bsn*$^{DAT}$cKO and *DAT*$^{Cre/+}$ mice displayed similar performances in terms of duration or latency to fall (Supplementary Figure 1H; t(13)=0.1266, p=0.9012, Student's t-test) and the speed they sustained (t(13)=0.0923, p=0.9278; data not shown) on accelerating rotarod.

Finally, *Bsn*$^{DAT}$cKO and *DAT*$^{Cre/+}$ mice were tested in the combined contextual and cued fear conditioning paradigm. Baseline freezing levels were unchanged between the genotypes (*Bsn*$^{DAT}$cKO: Freezing - 6.7 ± 1.2 %; *DAT*$^{Cre/+}$: Freezing - 6 ± 0.9 %;

t(13)=0.4651, p=0.6496; data not shown). While testing for the contextual fear memory, both the groups display similar freezing levels (Supplementary Figure 1I; t(13)=0.8974, p=0.3858, Student's t-test). 24 hrs later, mice were tested in neutral context to assess cued memory (CS) (Supplementary Figure 1J). Both groups exhibit similar and low freezing levels towards neutral tone (CS-) and profound freezing towards conditioned tone, CS+ (effect of test stimulus: F(1,13)=62.01, p<0.0001, two-way repeated measures ANOVA). However, no evidence was found for a genotype effect (F(1,13)=0.00, p=0.9657) or a genotype x test stimulus interaction (F(1,13)=0.00, p=0.9587). Together these data indicates that $Bsn^{DAT}$cKO mice did not display any fear memory alterations compared to $DAT^{Cre/+}$ mice.

It was previously showed that insertion of Cre under 3´UTR region of the *DAT* gene resulted in reduced expression of DAT protein in the striatum of homozygous and heterozygous mice compared to wild-type mice (Bäckman et al., 2006). So, to control for genetic background, $DAT^{+/+}$ (without Cre) and $DAT^{Cre/+}$ (heterozygous Cre) mice were tested in similar behavioral paradigms (Supplementary Figure 2) used for the characterization of $Bsn^{DAT}$cKO mice. Except for motor functions in rotarod paradigm (Supplementary Figure 2D, E), none of the other analyzed parameters was different between the $DAT^{+/+}$ and $DAT^{Cre/+}$ mice.

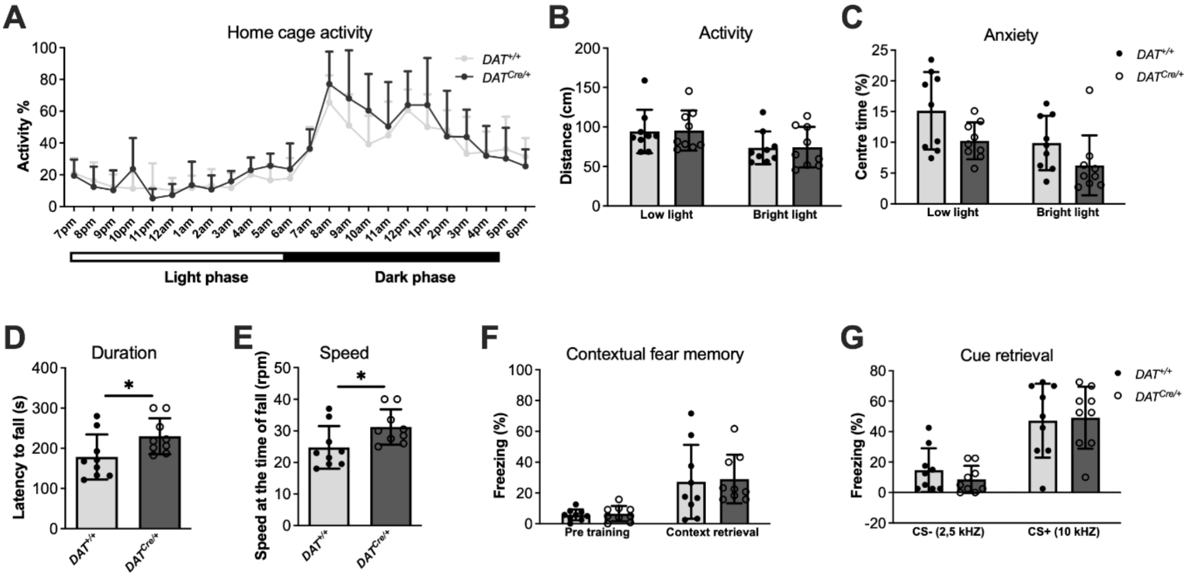


**Supplementary figure 2: Behavioral characterization of $DAT^{Cre/+}$ driver mice.** (A). Home cage activity monitoring, both $DAT^{+/+}$ and $DAT^{Cre/+}$ mice displayed normal circadian activity pattern [F(23,345)=31.55, p<0.0001, two-way repeated measures ANOVA] and no significant genotype effect [F(1,15)=0.68, p=0.4210] was observed. (B). Open field analysis of activity (distance travelled) during low light and bright light test sessions revealed significant effects of the test session [F(1,16)=103.62, p<0.0001, two-way repeated measures ANOVA] without any genotype effect [F(1,16)=0.01, p=0.9274] or effect of genotype x session interaction [F(1,16)=0.01, p=0.9358]. (C). Furthermore, as a measure of anxiety, the time spent by both groups at the center of the open field arena was analyzed. There was a significant effect of the test session [F(1,16)=19.01, p=0.0005] as well as main effect of genotype [F(1,16)=4.55, p=0.0487]. However, Bonferroni posttest does not reveal any significance differences. Statistical analysis using Student's t-test further revealed non-significant decrease in anxiety of $DAT^{Cre/+}$ mice during low light conditions [t(16)=2.107, p=0.0512]. (D, E). During analysis of motor functions using rotarod test, $DAT^{Cre/+}$ mice displayed superior performance in terms of duration or latency to fall [t(16)=2.162, p=0.0461, Student's t-test] and the speed they sustained [t(16)=2.209, p=0.0421] on accelerating rotarod. (F). During fear memory testing, both $DAT^{+/+}$ and $DAT^{Cre/+}$ displayed comparable pre-training baseline freezing levels, exhibited increased and comparable fear responses during contextual fear retrieval [Session effect: F(1,16)=25.72, p=0.0001] but no genotype effect [F(1,16)=0.06, p=0.8129] or genotype x session interaction [F(1,16)=0.01, p=0.9165]. (G). During memory towards auditory cue in a neutral context, both the genotypes showed low freezing levels to the non-conditioned auditory test stimulus (CS-), but strong freezing to the CS+ resulting in significant effect of test stimulus [F(1,16)=41.39, p<0.0001]. There was again no effect of genotype [F(1,16)=0.11, p=0.7445] or genotype x test stimulus interaction [F(1,16)=0.50, p=0.4883] observed. All values are mean ± SEM. $DAT^{+/+}$: N=9 and $DAT^{Cre/+}$: N=8; Data represented as mean ± SEM.
